# Pathophysiological response in experimental trauma-related acute kidney injury

**DOI:** 10.1101/2024.08.02.606294

**Authors:** Rebecca Halbgebauer, Lorena Schult, Onno Borgel, Arne Maes, Florian Weißhaupt, Christina Rastner, Alitsia Ast, Ludmila Lupu, Annette Palmer, Ulrich Wachter, Stefan A. Schmidt, Peter Boor, Reinhild Rösler, Sebastian Wiese, Greet Kerckhofs, Markus S. Huber-Lang

**Author notes:** Equally contributing senior authors (Greet Kerckhofs, Markus S. Huber-Lang). **Corresponding Author:** Markus Huber-Lang, M.D. Professor and Director Institute of Clinical and Experimental Trauma Immunology University Hospital of Ulm Helmholtzstr. 8/1 89081 Ulm Germany. ***Competing interests*** The authors declare that they have no competing interests.

## Abstract

**Background:** Trauma and shock often severely affect the kidneys. This can lead to trauma-related acute kidney injury (TRAKI), which significantly increases the risk of adverse outcomes.

**Methods:** To study the pathophysiology of TRAKI, we developed a murine model of combined blunt thoracic trauma and pressure-controlled hemorrhage that induces mild transient TRAKI.

**Results:** The mice showed early and transient increased plasma creatinine, urea, NGAL, and urine albumin, resolving 5 days after TRAKI induction. Despite normal kidney morphology, significant damage to proximal tubular cells and a loss of the brush border was observed. This included kidney stress responses, e.g., with induced heme oxygenase-1 expression in tubules. The upregulation of inflammatory mediators and kidney injury markers was followed by elevated leukocyte numbers, mainly consisting of monocytes/macrophages. Proteomic analyses revealed a distinct time course of intrarenal processes after trauma. 3D x-ray-based whole-organ histology by contrast-enhanced microcomputed tomography showed significant impairment of capillary blood flow, especially during the first day post THS, which was partly resolved by day 5.

**Conclusions:** Our novel model of murine TRAKI has revealed previously unknown aspects of the complex temporal pathophysiological response of the kidney along the nephron after trauma and hemorrhage, which may provide mechanistic starting points for future therapeutic approaches.

## Introduction

Physical trauma is a major global health issue caused by accidents, natural disasters and increasingly civil and military violence, and, notably, surgical interventions. Major posttraumatic complications include bleeding, infections, barrier and organ dysfunction (1), with frequent development of acute kidney injury (AKI) (2). The underlying etiology for AKI is manifold, including ischemia-reperfusion injury, hemorrhagic shock (3), crush syndrome (4), nephrotoxic drugs and contrast agents, major surgery (5), and sepsis (6).

In trauma-related AKI (TRAKI), all these triggers or their combination can induce a complex immuno-pathophysiological response (7), which, however, remains poorly defined. In the clinical setting, TRAKI often appears to be only temporary. Nevertheless, a recent big-data analysis in femur-fractured trauma patients indicated that even if patients with peritraumatic TRAKI were discharged with full renal recovery, the mortality rate 6 months post injury was more than 3 times higher than in patients without TRAKI. Moreover, the corresponding mortality rate correlated with the temporary Kidney Disease: Improving Global Outcomes (KDIGO) stage post trauma (8).

In a cohort of critically ill patients, the presence and recovery of AKI revealed even long-term mortality effects up to 15 years. Specifically, a subgroup with cardiac surgery-associated AKI who fully recovered during hospital stay exhibited mortality rates of 45% versus 30% for patients without in-hospital AKI (9). These facts underscore the need for deeper insights into underlying pathophysiological changes beyond the early period after trauma and their dynamic structure-function impacts.

At the molecular level, trauma results in the generation and release of multiple damage-associated molecular patterns (DAMPs) (10), such as heme (11), histones, and mitochondrial debris (12). These DAMPs can evoke a local renal and systemic inflammatory response, cellular damage and dysfunctional regeneration within the kidneys and intercommunicating organs (7, 13). After trauma, the reduction or loss of urine output as a key KDIGO criteria for AKI must be interpreted with utmost care. A small urine output could be caused by severe tissue damage or represent an appropriate physiological kidney adaptation to protect the trauma patient from hemodynamic instability (14).

Along the clearance trajectory from the afferent blood vessels via the glomerular filter and tubular system to the urine output, many structure-function-adaptive reactions or corresponding problems may occur after trauma. However, the pathophysiological focus in the context of trauma has traditionally been set on prerenal failure due to hemodynamic supply problems (e.g., shock) and hypoxia/ischemia-caused injury to the proximal tubules (e.g., acute tubular necrosis) (15), which seems to be, at least in part, a misconception (16). So far, changes beyond the initial rescue phase are rarely systematically investigated in the trauma context since biopsies in the peritraumatic and recovery phases could endanger the patient and raise ethical concerns.

Finally, most experimental AKI studies employ ischemia-reperfusion injury, transplantation, sepsis, or intoxication models. In contrast, our current study utilizes a retranslation model to simulate a remote, traumatic impact (blunt chest trauma) in combination with hemorrhagic shock and resuscitation. Our observation period extends up to 5 days post-injury, effectively reflecting a clinically transient TRAKI. This model, together with corresponding transcriptomics, proteomics, and 3D x-ray-based histology (17) of the kidneys, allowed us to spatially and temporally resolve the renal clearance trajectory and correlate our findings with established monitoring parameters. Our study thereby provides novel insights into dynamic changes of the renal blood supply and major molecular pathomechanisms in a clinically relevant model of TRAKI.

## Methods

### Animals

10-14-week-old male C57BL/6J mice with a mean body weight of 25.4 ± 0.3 g were purchased from Janvier Labs (Saint Berthevin, France) and had access to food and water *ad libitum.* The study was approved by the local University Animal Care Committee and the Federal Authorities for animal research, Tuebingen, Germany (approval 1409) and conducted according to the NIH Guide for the Care and Use of Laboratory Animals (2011).

### Trauma and hemorrhage in mice

Mice were randomly assigned to thoracic trauma and hemorrhage or sham treatment (n=8 each, as determined by power analysis). Analgesia was given by s.c. injection of 0.05 mg/kg body weight buprenorphine (Movianto, Neunkirchen, Germany) 30 min prior to trauma induction. Animals were anesthetized with 3% sevoflurane (Sevorane, Abbott, Wiesbaden, Germany) in oxygen. After shaving of the thorax, abdomen, and left hindleg, blunt thoracic trauma was applied: A blast cylinder was positioned centrally at 1.6 cm distance from the sternum, with the lower edge of the blast cylinder aligned with the lower osseous edge of the sternum. The chest was exposed to a defined blast wave with a pressure of 0.83 ± 0.27 bar for a duration of 1 – 2 ms. The blast wave induces bilateral lung contusion including intra-organ hemorrhage and edema formation without any damage to bony structures, a short period of apnea, and a transient decrease in heart rate and mean arterial pressure (18–20). After the trauma procedure, animals were placed on a heating pad with a rectal temperature probe and body temperature was kept at 37°C by a feedback loop.

For induction of severe hemorrhage and monitoring of the blood pressure and heart rate, the left distal *Arteria femoralis* was cannulated aseptically using polyethylene tubes (Smith, Grasbrunn, Germany). Pressure-controlled hemorrhage was induced by drawing blood for 5 – 10 min until a mean arterial pressure (MAP) of 30 ± 5 mmHg was reached; withdrawn blood was anticoagulated with heparin. The MAP of 30 ± 5 mmHg was maintained for 60 min by rebleeding if indicated by pressure measurement. After 60 min, the heparinized own blood was reperfused for 10 min. In the following, the animals were monitored for 60 min, and 2 *x* 500 µl Jonosteril (Servoprax, Wesel, Germany) was given i.v. for fluid substitution. Afterwards, the artery was closed aseptically by applying 5/0 Prolene sutures (Johnson & Johnson, New Brunswick, NJ, USA) and animals were allowed to wake up. 0.05 mg/kg body weight buprenorphine was injected s.c. every 8 h during the first 24 h after trauma; afterwards, analgesia was given when necessary. Sham animals received analgesia and were anesthetized for 2 h; a small skin incision was applied in the left hindleg and sutured after 2 h. 4 h, 24 h, and 5 d after trauma or sham treatment, the experiment was terminated by cardiac puncture under deep anesthesia. Kidneys were explanted and processed for histological evaluation, electron microscopy analysis, contrast-enhanced microfocus x-ray computed tomography (CECT) or snap-frozen in liquid nitrogen. Heparinized blood was collected by cardiac puncture, centrifuged at 800 x g and 4 °C for 5 min, followed by centrifugation at 13000 x g and 4 °C for 2 min, and plasma was stored at - 80°C. Urine was collected by bladder puncture, centrifuged at 800 x g and 4 °C for 5 min, and urine samples were stored at -80 °C until analysis. Researchers were blinded to sample group allocation for all subsequent analyses. Whenever samples were available for each animal, they were included in the analysis.

### Plasma and urine analysis

Plasma und urine NGAL as well as urine albumin were determined using commercial ELISA kits (Mouse NGAL ELISA Kit, LifeSpan BioSciences, Seattle, USA; Albumin Simple Step ELISA Kit, Abcam). Results were normalized to plasma protein content as a corrective measure for fluids given. Plasma creatinine and urea were measured as described elsewhere (21).

### Classical 2D histology

Tissue samples were fixed in 3.7% formaldehyde in PBS (Fischar, Saarbruecken, Germany) for 24 h, dehydrated, and embedded in paraffin. 3-4 µm sections were cut using a microtome, followed by rehydration in a graded alcohol series. For histological evaluation, periodic acid– Schiff staining was performed using a commercial kit (Merck, Darmstadt, Germany). Images were taken using an Axio Imager M1 microscope and the ZEN 3.0 software (Zeiss, Oberkochen, Germany).

### Immunohistochemistry and immunofluorescence

For immunohistochemistry and -fluorescence, slides were blocked with 5% normal goat serum. After antigen retrieval, tissues were incubated with anti-heme oxygenase-1 antibody (1:100, clone EP1391Y, Abcam, Cambridge, UK), anti-fibrinogen beta chain antibody (1:8000, clone EPR18145-84, Abcam), anti-CD45 antibody (1:500, clone EPR20033, Abcam), anti-F4/80 antibody (1:100, clone Cl:A3-1, Bio-rad, Feldkirchen, Germany), and anti-CD10 antibody (1:300, clone EPR22867-118, Abcam). For visualization of heme-oxygenase 1, fibrinogen beta chain, and CD45, the Dako REAL detection Alkaline phosphatase Kit (Dako, Glostrup, Denmark) was used. For detection of F4/80, slides were incubated with a peroxidase goat-anti-rabbit antibody (Jackson ImmunoResearch, Cambridge, UK), followed by development using DAB substrate (Merck). For assessment of CD10 staining, an Alexa-Fluor 488-goat anti-rabbit IgG (Thermo Fisher, Waltham, MA, USA) was used. Images of five randomly selected regions were taken in even distribution over the entire cortex in 100-fold magnification using an Axio Imager M1 microscope and the ZEN 3.0 software (Zeiss). Cells positive for CD45 staining per field of view as well as the percentage of CD10-positive tubules were counted in a blinded fashion using the “Cell Counter” plug-in for ImageJ (NIH, USA).

### TUNEL assay

For terminal deoxynucleotidyl transferase dUTP nick end labeling (TUNEL) staining, slides were stained using the In-Situ Cell Death Detection Kit, Fluorescein (Merck) following the manufacturer’s instructions. Images were taken using an Axio Imager M1 microscope and the ZEN 3.0 software (Zeiss).

### Transmission electron microscopy (TEM)

Cortical tissue was fixed with 2% paraformaldehyde and 2.5% glutaraldehyde (Fluka/Fisher Scientific, Schwerte, Germany) containing 1% saccharose (Roche, Switzerland) in phosphate buffer (pH 7.3), washed 5 times with 0.01 M PBS buffer and post-fixed in 2% aqueous osmium tetroxide (Fluka). Samples were then dehydrated in a graded series of 1-propanol, block stained in 1% of uranyl acetate and embedded in Epon (Fluka). Ultra-thin sections (80 nm) were contrasted with 0.3% lead citrate for 1 min and imaged in a JEM-1400 (Jeol, Freising, Germany) in 8000x magnification.

### Quantitative real-time PCR and microarray analysis

RNA was isolated from snap-frozen mouse kidneys using the RNeasy Mini Kit (Qiagen, Hilden, Germany). Reverse transcription was performed using AffinityScript qPCR cDNA Synthesis Kits (Agilent Technologies, Waldbronn, Germany) and RT-qPCR was carried out employing the Brilliant III Ultra-Fast SYBR® Green Master Mix (Agilent Technologies). Primers are listed in **Supplemental file 1, Table S1**. Fold expression in comparison to respective sham animals was calculated using the 2^-ddCT^ method.

For microarray analysis, only samples with an RNA integrity number ≥ 9.1 were used. Per sample, 200 ng of total RNA and 5.5 μg single-stranded DNA were applied using a GeneChip Fluidics Station 450 (Affymetrix, Santa Clara, USA). After hybridization of single-stranded DNA to Mouse Gene 1.0 ST GeneChip Arrays, arrays were scanned by a GeneChip scanner 3000 (both Affymetrix). Images were analyzed using Affymetrix Expression Console Software and BRB-ArrayTools. After applying the robust multiarray average method, normalized values of raw feature data and log2 intensity expression summary values were computed.

### Contrast-enhanced microfocus x-ray computed tomography (CECT)

For CECT analysis of the kidneys, blood vessels of the left kidney from *n* = 5 animals per group were ligated before cardiac exsanguination in order to keep the blood volume *in situ*. Kidneys were fixed in 4% formaldehyde for 24 h, washed in PBS and stained in 35 mg/ml 1:2 hafnium(IV)-substituted Wells-Dawson polyoxometalate (Hf-WD POM) in PBS for 10 days at room temperature while gently shaking. Prior to image acquisition, kidneys were wrapped in parafilm. The upper pole was imaged using a Phoenix NanoTom M (GE Measurement and Control Solutions, Germany) with a 180 kV/15 W energy X-ray tube source and employing a diamond-coated tungsten target (for settings, refer to **Supplemental file 1, Table S2**). The Datos|x software (GE Measurement and Control Solutions) was used to reconstruct datasets with the beam hardening correction (value: 8), inline median filter and ROI-CT filter active. Reconstructed and exported 16-bit slices (.tiff) were converted to 8-bit slices (.bmp) with an in-house developed MATLAB script, whilst simultaneously windowing the histogram range to the dynamic range of the dataset (22). Datasets were visualized in 3D and segmented using Avizo 2022.1 software (Thermo Fisher Scientific, Bordeaux, France). For normalization between samples, employing Avizo’s module “interactive thresholding”, a grey value threshold range was selected to include >90% of the cross-section of a central *Arteria arcuata renis.* Four volumes of interest (VOIs) per kidney at 1 mm^3^ each were selected in the cortical region and segmented using the “watershed” tool. An example of the filtering process is shown in **Supplemental file 1, Fig. S1**. Data was converted and assessed using Bruker CTan 1.18.9.0 (Blue Scientific Limited, Cambridge, UK). Using “Thickness analysis”, distribution in vessel diameter was assessed, and relative volumes for capillaries (2-10 µm) and arterioles/venules (10-30 µm) were measured.

### Mass spectrometric analysis

For tandem mass tag (TMT, Thermo Scientific) six-plex labelling, 100 µg of sample were labelled using TMT according to the manufacturers protocol. Mass spectrometric characterization was performed as published previously (23). In brief, following equal mixing, combined samples were fractionated using strong cation exchange (SCX) chromatography on a BioRSLC (Thermo Scientific). After desalting and vacuum drying, samples were reconstituted and mass-spectrometrically analyzed on an Orbitrap Elite instrument (Thermo Scientific).

Database searches were performed using MaxQuant Ver. 1.6.3.4 (www.maxquant.org) (24). For peptide identification and quantitation, MS/MS spectra were correlated with the UniProt mouse reference proteome set (www.uniprot.org), employing the build-in Andromeda search engine (25). The respective TMT modifications and carbamidomethylated cysteine were considered as a fixed modification along with oxidation(M), and acetylated protein N-termini as variable modifications. False discovery rates on both peptide and protein level were set to 0.01.

Subsequent data analysis was performed employing MS Excel and GraphPad Prism. For statistical analysis, a combination of t-tests and outlier analysis based on significance B was used. For the latter and for hierarchical clustering, the Perseus software suite (https://maxquant.org/perseus/) was employed with standard parameters. Significance levels for tests were set to 0.05. Proteins having passed both tests as being significantly regulated were classified as category I, while proteins considered as outliers using significance B were classified as category II. Pathway and process enrichment analysis was performed using www.string-db.org (26).

### Statistical analysis

GraphPad Prism v. 9 (Graphpad Software, Inc., USA) was used for statistical assessment. Data is shown as mean; error bars indicate the standard error of the mean. Outliers were identified employing Prism’s ROUT method. For comparisons between THS and corresponding sham animals at respective time points, Mann-Whitney-U test was used. Data in three or more groups was compared using Kruskal-Wallis one-way ANOVA on ranks with Dunn’s post-hoc testing. A p-value below 0.05 was considered significant.

## Results

### Transient TRAKI after murine trauma and hemorrhage

In this study, a mouse model of combined thoracic trauma and pressure-controlled hemorrhagic shock (THS) was established. The study protocol is shown in **Fig. 1A**. In THS animals, the MAP was reduced to 30 mmHg for 60 min, which was accompanied by a compensatory increase in the heart rate (**Fig. 1B**). Urine albumin (**Fig. 1C**) as well as plasma creatinine, urea and NGAL (**Fig. 1D-F**) concentrations were significantly elevated after THS especially during the early posttraumatic phase.

**Fig. 1:**
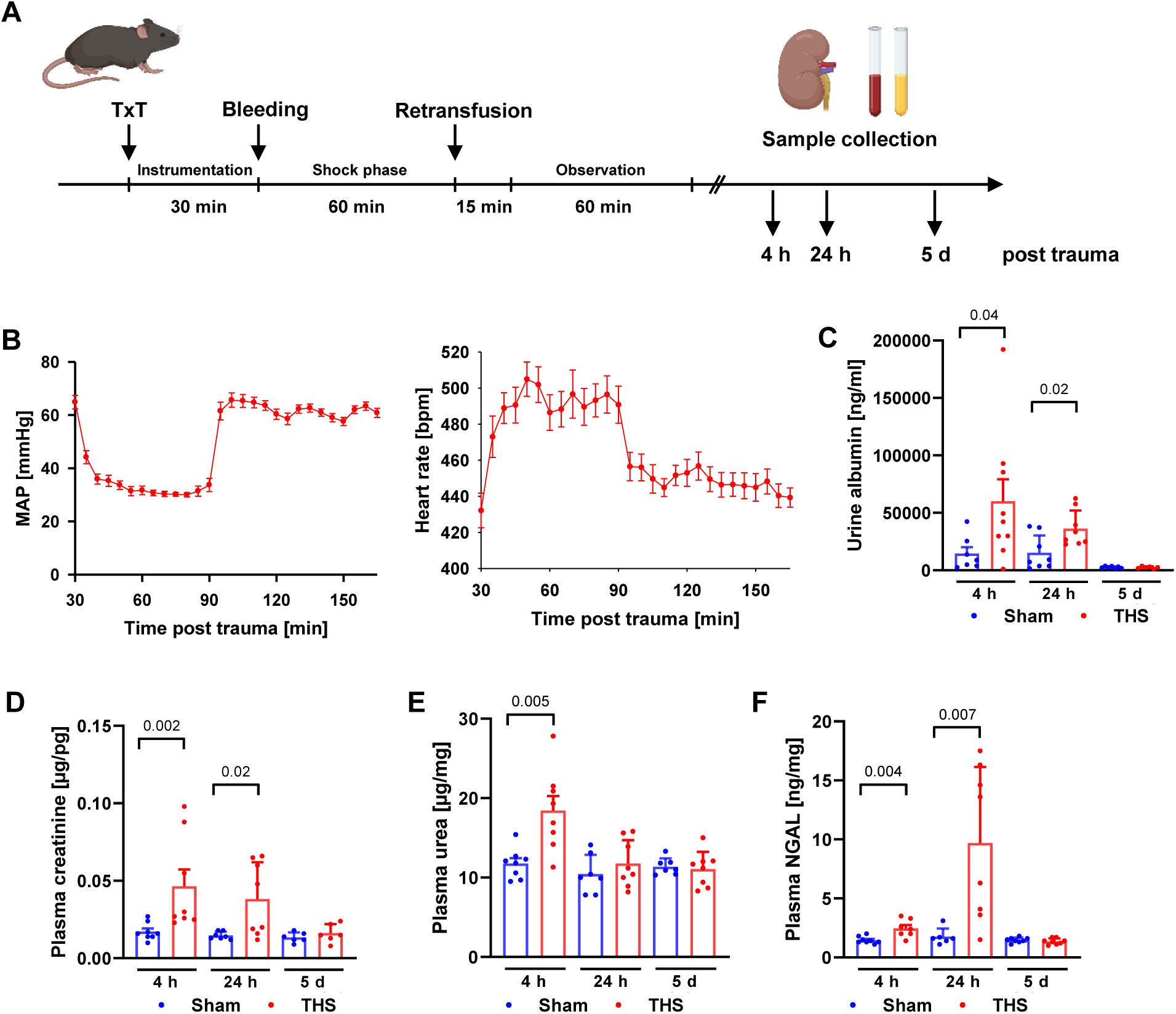
A novel mouse model of blunt chest trauma and hemorrhage results in mild transient TRAKI. **A,** Experimental design: Mice underwent thoracic trauma, followed by pressure-controlled shock (THS) for 60 min at 30 ± 5 mmHg. After retransfusion with the drawn blood, animals were monitored and allowed to wake up. Kidneys, blood, and urine were collected 4 h, 24 h, and 5 d after trauma. **B**, Mean arterial pressure (MAP) and a compensatory increase in the heart rate in THS animals. **C**, Albumin urine concentrations after sham treament or THS. **D-F**, Creatinine (**D**), urea (**E**), and neutrophil gelatinase-associated lipocalin (NGAL) (**F**) in plasma from mice after THS/sham treatment. Graphs indicate the mean with SEM from *n* = 26-27 animals (**B**,**C**) or *n* = 6-8 animals per group (**D**-**H**). Comparisons of THS animals with corresponding sham groups were performed using Mann-Whitney-U test.

### Unaltered glomerular structure and proximal tubular injury after trauma

Overall kidney morphology as assessed by classical 2D histology remained normal in animals after THS (**Fig. 2A**). Transmission electron microscopy (TEM) (**Fig. 2B**) and gene expression analysis (**Fig. 2C**) did not reveal any pathological changes in podocyte morphology or functional marker expression. However, we observed proximal tubule damage as reflected by a loss of the brush border (**Fig. 2D,F,G**) and increased expression of proximal tubule injury markers (**Fig. 2E**). Distal tubular cells were not affected by THS as assessed by TEM (**Supplemental file 1, Fig. S2**).

**Fig. 2:**
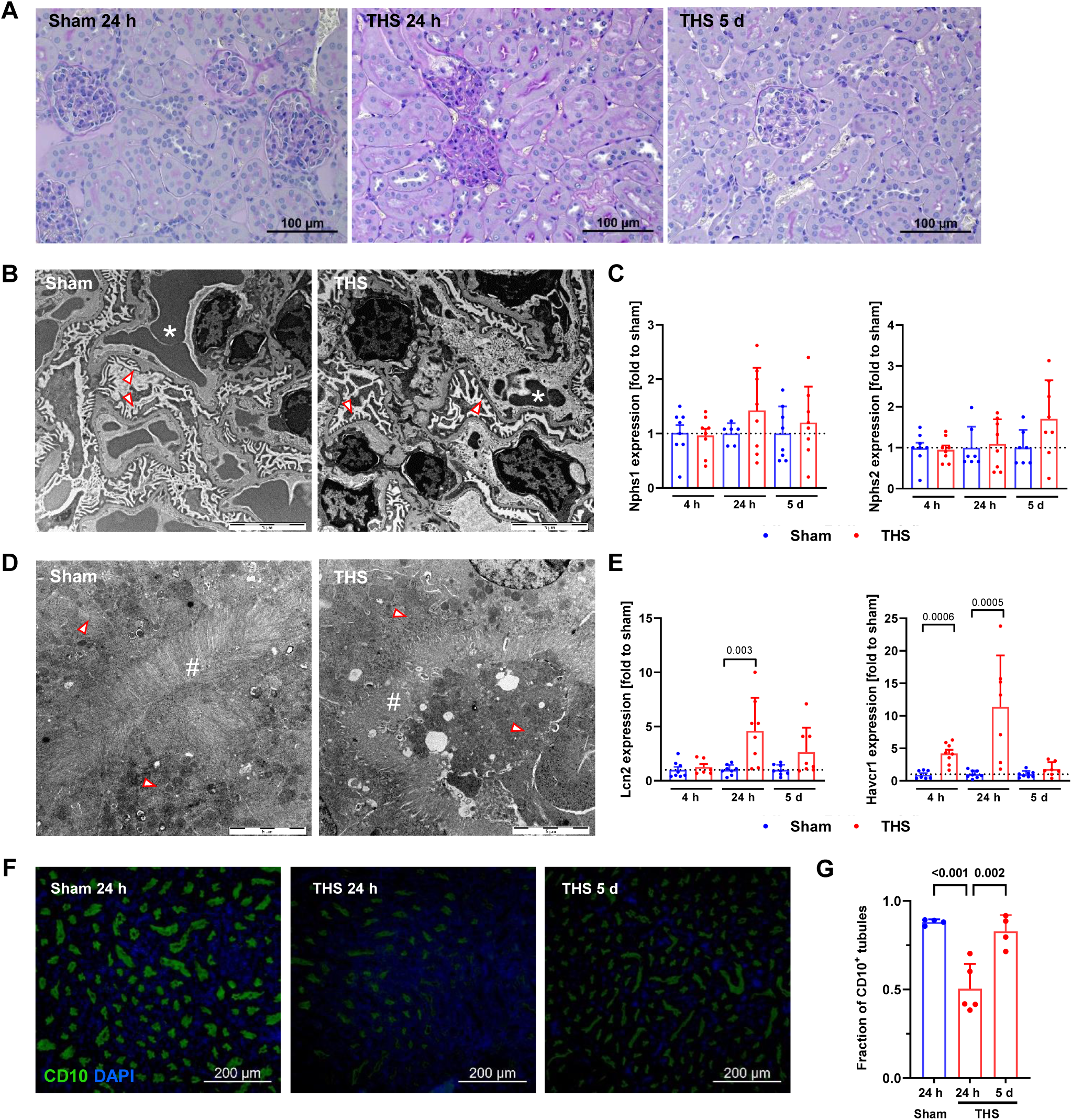
Glomerular structure and proximal tubular injury after trauma. **A**, Normal tissue morphology as assessed by PAS staining. Images are representative of *n* = 5 animals per group. **B**, Transmission electron microscopy (TEM) analysis of the glomerulus with unaltered podocyte morphology 24 h after THS or sham treatment. Images are representative of *n* = 5 animals per group. Star symbols indicate erythrocytes in glomerular capillaries, arrowheads indicate podocyte pedicles along the blood-urine barrier. **C**, Tissue expression of the podocyte functional proteins nephrin (Nphs1) and podocin (Nphs2) during the time course after THS in comparison to control animals. **D**, Transmission electron microscopy (TEM) analysis of the proximal tubulus with less regular brush border structure 24 h after THS in comparison to sham treatment. Images are representative of *n* = 5 animals per group. Hash symbols indicate brush border; arrowheads indicate tubule cell cytoplasm. **E**, Tissue expression of the proximal tubule injury markers neutrophil gelatin-associated lipocalin (NGAL) and kidney injury molecule-1 (KIM-1) 24 h after THS or sham treatment. **F**, CD10 staining of sham and THS animals at indicated timepoints visualizing the brush border. Images are representative of *n* = 5 animals per group. **G**, Fraction of CD10^+^ tubules per field of view (100x magnification) from *n* = 5 animals. Scale bars are indicative of 5 µm (B,D).

### Pathophysiological mechanisms of TRAKI

In a bulk gene expression screening, we identified 25 significantly altered genes in kidneys 24 h after THS compared to sham animals (**Supplemental file 1, Fig. S3A, Source data 1**). The upregulation of Hmox1 was confirmed on a protein level (**Fig. 3A**) and verified by qPCR (**Fig. 3B**). An increase in renal IL-34 expression as verified by qPCR (**Fig. 3C**) resulted in a delayed tendential, but non-significant increase in leukocyte numbers (**Fig. 3D,E**). We next sought to characterize the cell type and found a tendency of reduced F4/80^+^ cell numbers 24 h and a slight, but non-significant elevation of monocyte marker expression in kidney tissues 5 d after THS (**Supplemental file 1, Fig. S3C,D**) while numbers of CD3^+^, CD19^+^, and Ly6G^+^ cells remained unaltered (data not shown). Since several genes identified were associated with apoptosis, we performed TUNEL staining. However, THS did not induce an increase in the number of TUNEL-positive cells compared to sham-treated animals (**Supplemental file 1, Fig. S3B**). In order to assess renal blood filling, we performed CECT and analyzed the volume of filled vessels (**Fig. 3F,G**). In comparison to controls with a heterogeneous distribution of capillary filling in volumes of interest distributed over renal cortices, THS animals demonstrated a significantly reduced capillary blood filling on day 1, which partly normalized until day 5; a similar trend was visible for arterioles and venules (**Fig. 3H**).

**Fig. 3:**
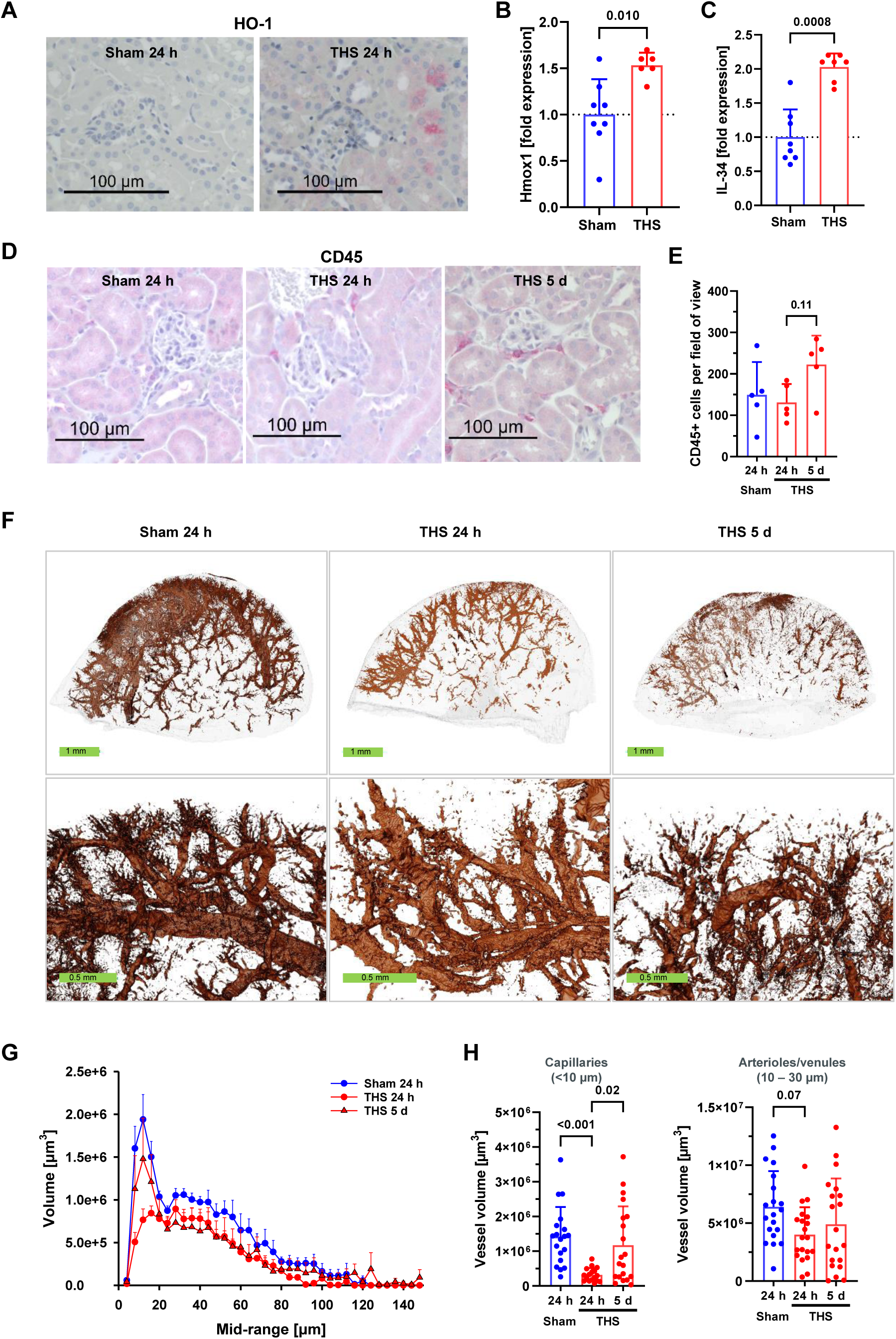
Intrarenal stress responses and reduced perfusion after trauma and hemorrhage. **A,** Immunohistochemical staining of the stress response factor heme oxygenase-1 (HO-1) in kidneys from mice 24 h after THS or sham treatment. HO-1 is stained in red; nuclei are blue. Images are representative of at least *n* = 5 animals per group. **B**, Gene expression of HO-1 (gene: Hmox1) in kidneys from mice 24 h after THS or sham treatment (*n* = 7-8). **C**, Renal IL-34 gene expression 24 h after THS or sham treatment; *n* = 7-8. **D**, CD45 staining in kidneys at the indicated time points after THS or sham treatment. Leukocytes are stained in red, nuclei in blue. Images are representative of at least *n* = 5 animals per group. **E**, CD45^+^ cells per field of view in five randomly selected regions from *n* = 5 animals per group. **F**, CECT-based 3D rendering of the blood vessel network in the upper kidney poles at the indicated time points after THS or sham treatment in overview (upper panel) and in detail (lower panel). Images are representative of *n* = 5 animals per group. **G**, Vessel volume distribution according to the mid-range vessel diameter for kidneys explanted 24 h or 5 d after sham treatment or THS. **H**, Vessel volume for capillaries (2-10 µm diameter) and arterioles/venules (10-30 µm diameter) in kidneys at the indicated time points after THS or sham treatment.

In a proteomic approach, we aimed to assess the alterations in renal protein abundance to gain more insight in TRAKI pathophysiology on a protein level. We found a distinct course of proteomic alterations depending on the time point post THS (**Fig. 4A-C, Source data 2**). Overall, alterations in THS vs. sham animals were most prominent 24 hours post trauma (**Fig. 4D**); a subset of proteins demonstrated a linear increase or decrease in the THS/sham ratios during the observation period **(Fig. 4E)**. The acute posttraumatic phase especially affected pathways involved in renal tubular secretion and ribosomal function (**Fig. 4A,F**). Here, we detected a ∼20% loss in ATPase Na+/K+ Transporting Subunit Alpha 1; however, classical 2D histological staining did not reveal changes in Na+/K+ ATPase tissue expression or distribution (**Supplemental file 1, Fig. S4A**). 24 hours post THS, we found changes predominantly in coagulation processes and increased abundance of fibrin components (**Fig. 4B,G**), which was confirmed by immunostaining of fibrin (**Supplemental file 1, Fig. S4B**). On day 5, we did not detect alterations in any of the pathways relevant during the acute to intermediate phase after trauma; however, some signs of epigenetic remodeling were apparent (**Fig. 4C,H**).

**Fig. 4:**
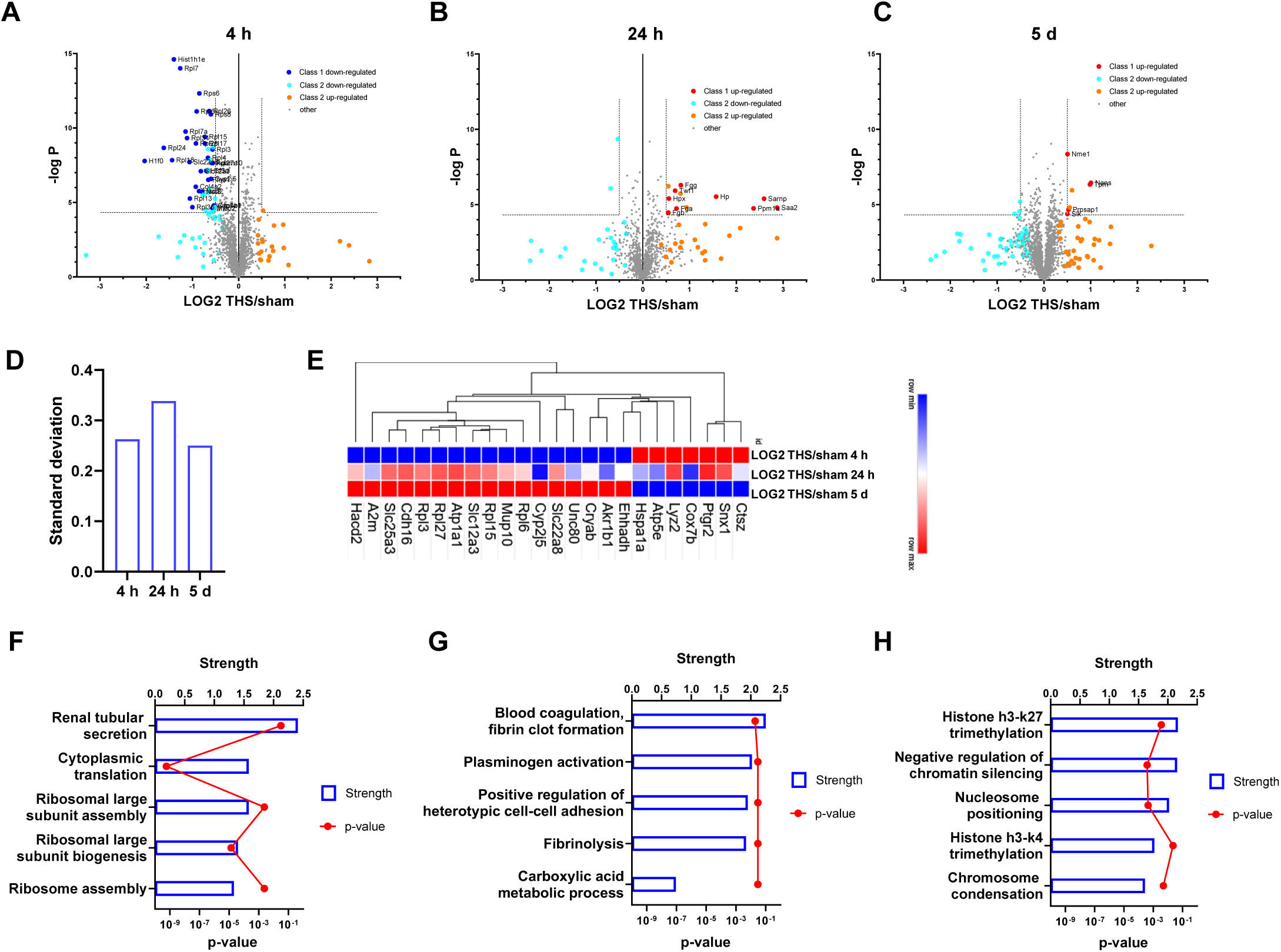
Temporal resolution of potential pathomechanisms behind TRAKI. **A**-**C,** Altered protein abundance in bulk kidney tissue 4 h (**A**), 24 h (**B**), and 5 d (**C**) after trauma and hemorrhage. Volcano plots show protein ratios in THS compared to sham animals, with protein allocation to class 1 and 2 based on t-test p < 0.05 (*n* = 3 animals, respectively). **D**, Overall standard deviation of protein ratios observed in the proteomic experiments demonstrating the highest degree of expression shifts 24 h after THS. **E**, Heatmap and dendrogram of protein abundance ratios of THS compared to sham animals with linear increase or decrease during the period of observation at the indicated time points based on a hierarchical clustering of a selection of proteins. **F-G**, Pathway analysis of proteins found to be statistically significant (non-grey in A-C) 4 h (**F**), 24 h (**G**), and 5 d (**H**) after THS compared to respective sham animals using string-DB.

## Discussion

In the present novel TRAKI mouse model, induced by remote blunt chest trauma and severe hemorrhagic shock to reflect a common combinatory injury pattern, followed by a hemodynamic resuscitation period within a clinically relevant time axis, we observed morphological and biochemical signs of temporary TRAKI. The plasma retention of urinary substances, the appearance of renal damage markers, and the development of albuminuria all indicate trauma-induced changes beyond the renal functional reserve, defining an AKI 1B stage according to the concept proposed by the Acute Disease Quality Initiative (ADQI) (27). The manifestation of TRAKI was especially pronounced 24 h after trauma, and the majority of indicators returned to baseline levels within 5 days. In search for underlying kidney changes and structure-function problems of TRAKI, this systematic investigation along the blood-urine clearance trajectory could uncover novel mechanistic aspects.

Starting at the renal *blood supply*, the hemodynamics of shock with a defined time-limited drop in MAP and a reflexively enhanced heart rate provided an established trigger for the development of TRAKI (7). In trauma patients, hemorrhagic shock is the major driver of TRAKI, which regularly develops within 5 days after trauma (3, 28, 29). However, unlike cases of transplantation or surgical cross-clamping where a no-flow phenomenon drives subsequent renal ischemia-reperfusion injury (IRI), TRAKI is rather characterized by a low-flow phenomenon (7), which should - in principle - be completely cleared upon resuscitation. Yet, in a sheep model of various degrees of hypoperfusion, the authors showed that reducing renal perfusion by 80% for 2 hours only resulted in temporary AKI which fully resolved rather rapidly (within 8 h) (30). It is therefore tempting to speculate that, in addition to severe hypoperfusion, traumatic tissue damage is required for the development of TRAKI. Indeed, in a pig model of 3 h of hemorrhagic shock plus minor surgical trauma (due to extended instrumentation), the renal blood flow was not restored even 24-48 hours after hemorrhagic shock despite a guideline-based resuscitation protocol (31). Furthermore, studying the renal perfusion in the present TRAKI model revealed a significant reduction of blood vessel filling 24 h after trauma/hemorrhagic shock, affecting primarily the capillaries (**Fig. 3E-G**). Importantly, these morphological signs of compromised perfusion persisted up to 5 days after resuscitation. The reasons for the reduced blood content can only be speculated upon, but it is likely that the trauma-induced sympatho-adrenergic drive results in a prolonged renal vasoconstriction irrespective of systemic vasotonic changes, as seen in a large animal study of THS (31). Another possible mechanism could be a rarefication of the peritubular capillaries, as previously shown in severe IRI models in mice weeks after the insult (32). However, although the rarefication pattern of the affected vessels (32) appeared roughly similar in our TRAKI model (despite using different imaging methods), such a rarefication would occur unexplainably early in our study. Theoretically, enhanced interstitial pressure surrounding the capillaries or intraluminal thrombi could also contribute to depressed perfusion. However, these speculations require further investigations, which could utilize the employed CECT x-ray-based 3D histology imaging technique with the potential of subsequent post-imaging classical 2D histological staining opportunities.

Concerning the *glomerular filter* as next spatial module, the appearance of proteins, including albumin, in the urine up to 24 h after trauma might, at first glance, suggest some temporary damage to the glomerular sieve. However, neither the classical 2D histological nor TEM analyses identified significant morphological changes (except some scattered irregularities in the thickness of the basal membrane in the trauma group). Damage to the foot processes of podocytes was excluded by electron microscopy and unaltered podocyte functional markers. In line, a previous report charged reactive oxygen species (which are also generated in trauma and hemorrhagic shock) for causing reversible proteinuria and glomerular filter defects without apparent structural abnormalities (33). Albuminuria might also be caused by reabsorption deficits in the downstream kidney epithelium of the tubular system, supported by our proteome profile indicating a loss of various transporter systems.

The *tubular system* was indeed massively affected in all animals after THS. Loss of the brush border of proximal tubules peaking at 24 h and partly recovering 5 days after trauma, TEM-signs of tubular epitheliopathy in the proximal (but not distal) tubule, and enhanced kidney damage markers such as NGAL and KIM-1, indicate severe tubular injury and resorptive dysfunction early in TRAKI. These are reflected by distinct proteomic changes and represent a vital danger to the electrolyte-water balance of the traumatized organism. Almost half a century ago, a theoretical consideration proposed the life-saving importance of glomeruli in case of a damaged tubular system: since the entire circulating plasma volume is filtered and reabsorbed twice an hour, the juxta-glomerular feedback loop needs to dramatically reduce the glomerular filtration rate; otherwise, systemic electrolytes and water would be lost within a few hours (14). Whereas the energy source for glomerular filtration is provided by the left ventricle and blood pressure, metabolism of the kidney cells provides the energy for resorptive processes (14). The reduced blood pressure in the present model – as normally observed in the clinical setting of trauma and shock – is rapidly reversed, whereas cellular metabolism might be altered for a longer time period (34). In turn, this imbalance might prolong the recovery of the kidneys post trauma, as demonstrated in our model, but also in clinical TRAKI (28).

Concerning kidney infiltration by *inflammatory cells*, a recent study proposed that the mode of AKI determines neutrophil infiltration; bilateral renal ischemia-reperfusion-injury (IRI), but not septic AKI, resulted in robust neutrophil recruitment (35). In our non-ischemic, non-septic TRAKI model, CD45^+^ leukocytes, consisting mainly of F4/80^+^ monocytes/macrophages, were detected in the kidneys after trauma, whereas neutrophils seemed to play, if at all, a minor role. In murine IRI and rhabdomyolysis models, augmented renal monocytes/macrophages (in the absence of macrophage migration inhibitory factor) have been proposed as the driving mechanism for tubular injury 24 h after the insult (36), supporting our findings. However, in our TRAKI model, the monocytes were not recruited in the early phase (at 24 h) but rather in the later phases after trauma (until day 5). Accordingly, IL-34, which is generated in tubular epithelial cells and promotes monocyte proliferation and survival, was upregulated in the kidneys 24 h after trauma and likely functioned as the driving force for the delayed monocyte increase. In murine IRI-AKI, IL-34 also fostered monocyte-caused kidney destruction and induced chronic kidney disease (CKD) (37), which in principle, could be the case in TRAKI as well, but was not further investigated as a potential therapeutic approach as a limitation of this study. Taken together, inflammatory cell recruitment in the case of TRAKI displayed its own distinct dynamic features, different from pure ischemic or septic AKI (35).

Addressing the dynamics on a *molecular level*, the proteome signature revealed novel concepts for TRAKI. As expected, the proteome analysis pointed to significant impairments in tubular absorption and secretion. We observed a significant reduction of the Na^+^-K^+^-ATPase (ATP1a1) and of various solute carriers, including sodium-glucose cotransporter (Slc5a2), sodium-neurotransmitter symporter (Slc6a19), and cation-chloride cotransporter (Slc12a3) early after trauma. Such mainly sodium-driven channelosome alterations impact electrolytes, water-osmoregulation, pH balance, and transport mechanisms. The enhanced myoglobin levels may reflect accumulation of muscle tissue debris within the kidneys, which also can function as DAMPs for the kidney response. Furthermore, detected alterations in ribosomal proteins may reflect induction and/or disturbances of early repair mechanisms. Of note, 24 hours after trauma, proteins involved in coagulation (such as fibrin) and synchronic fibrinolysis (plasminogen activation) appeared in TRAKI, indicating the induction of renal thromboinflammation. Whether the renal activation of the coagulation/fibrinolysis system also resulted in compromised capillary perfusion as observed in 3D-virtual analysis remains to be determined. Another mechanism concerns stress-induced heme oxygenase-1 (HO-1) which results in the clearance of free heme acting as a potent DAMP after trauma (38). HO-1 was slightly but significantly upregulated in the kidneys 24 hours after trauma, thus providing an early anti-oxidative, anti-inflammatory renoprotective mechanism (38). In later periods after TRAKI, when functional markers had largely normalized, epigenetic alterations such as significant decreases in histone H.1.0/2/3/4 and histone H4 were predominantly found. Moreover, carbonic anhydrase, which mainly acts within the proximal tubule and catalyzes the interconversion of CO_2_ and H_2_O, was significantly reduced in the kidneys 5 days after trauma. Centrally involved in maintenance of the acid-base balance and water resorption, the drop in carbonic anhydrase may help restore diuresis.

Another limitation of our study is that the animal protocol did not allow us to extend the duration of shock or to assess renal function and repair beyond day 5, thus precluding conclusions regarding the development of renal fibrosis and subsequent chronic renal dysfunction as observed by others after prolonged hemorrhage in mice (39). Future studies will determine whether trauma and hemorrhage are causative for sustained renal injury and evaluate potential therapeutic interventions to clarify a causal relationship between the proposed pathomechanisms and TRAKI. However, our work in a large population study in humans suggests that even transient renal dysfunction (with full functional recovery), e.g. during clinical treatment of major fractures, is highly associated with increased mortality over the next few months (8).

### Conclusions

The modern translation of acute renal “success” rather than “failure” (14) may characterize morphological, biochemical, and functional key changes as adaptive mechanisms of TRAKI striving to overcome tissue damage and preserve vital kidney function post trauma. However, the findings may also have important mechanistic implications for persistent kidney problems and dysfunctional kidney-organ crosstalk (7) resulting in a poor outcome post trauma. Therefore, the present insights into the temporal-spatial immune-pathophysiological response of TRAKI provides mechanistic starting points for therapeutic approaches that need to be addressed in future studies.

## Declarations

### Ethics approval and consent to participate

The study was approved by the local University Animal Care Committee and the Federal Authorities for animal research, Tuebingen, Germany (approval 1409) and conducted according to the NIH Guide for the Care and Use of Laboratory Animals (2011).

### Consent for publication

Not applicable.

### Availability of data and materials

The datasets supporting the conclusions of this article are included within the article and its additional files.

### Authors’ contributions

RH, AP, GK and MHL contributed to conception and design of the study. RH, LS, OB, AM, FW, CR, AA, LL, AP, and UW performed experiments and contributed to data collection. RH, PB, SS, and GK contributed to data analysis. MHL supervised the study. RR and SW performed the proteomic experiments as well as the accompanying data and related statistical analysis. RH and MHL performed the statistical analysis and wrote the first draft of the manuscript. All authors contributed to manuscript revision and approved the submitted version.

## Acknowledgements

We would like to express our gratitude to Sonja Braumueller, Bettina Berger, Lena Doerfer, and Anke Schultze (Ulm University Medical Center) for excellent technical assistance. We also thank Dr. Karl-Heinz Holzmann and Karin Lanz at the Core Facility Genomics (Ulm University Medical Center) and Prof. Dr. Paul Walther at the Core Facility Electron Microscopy (Ulm University Medical Center) for their expert support. We would like to thank Antoine Chretien (UCLouvain) for his help in dissecting the kidneys for the technique optimization experiments, and Pierre Schneidewind (UCLouvain) for the sectioning and colorimetric staining of the biological samples.

---

**Supplemental Figure 1.**
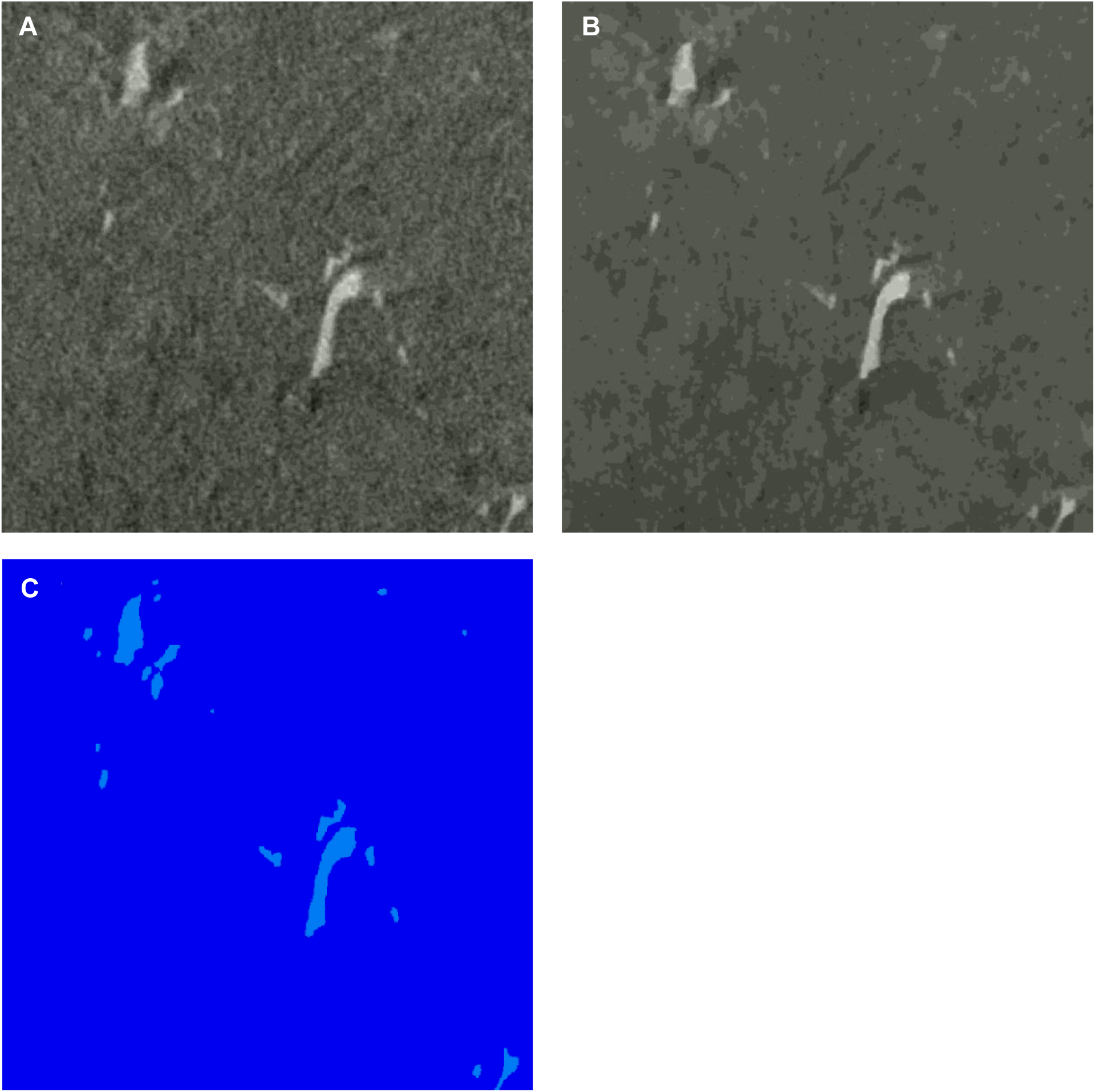
Processing of CECT images. **A**, raw image after subvolume extraction. Blood vessels are visible as light grey structures compared to the surrounding tissue. **B**, image after processing using “non-local-means filter” and “unsharp masking filter”. **C**, image after application of the “watershed” tool. Light blue areas correspond to blood vessels, dark blue areas to surrounding tissue.

**Supplemental Figure 2.**
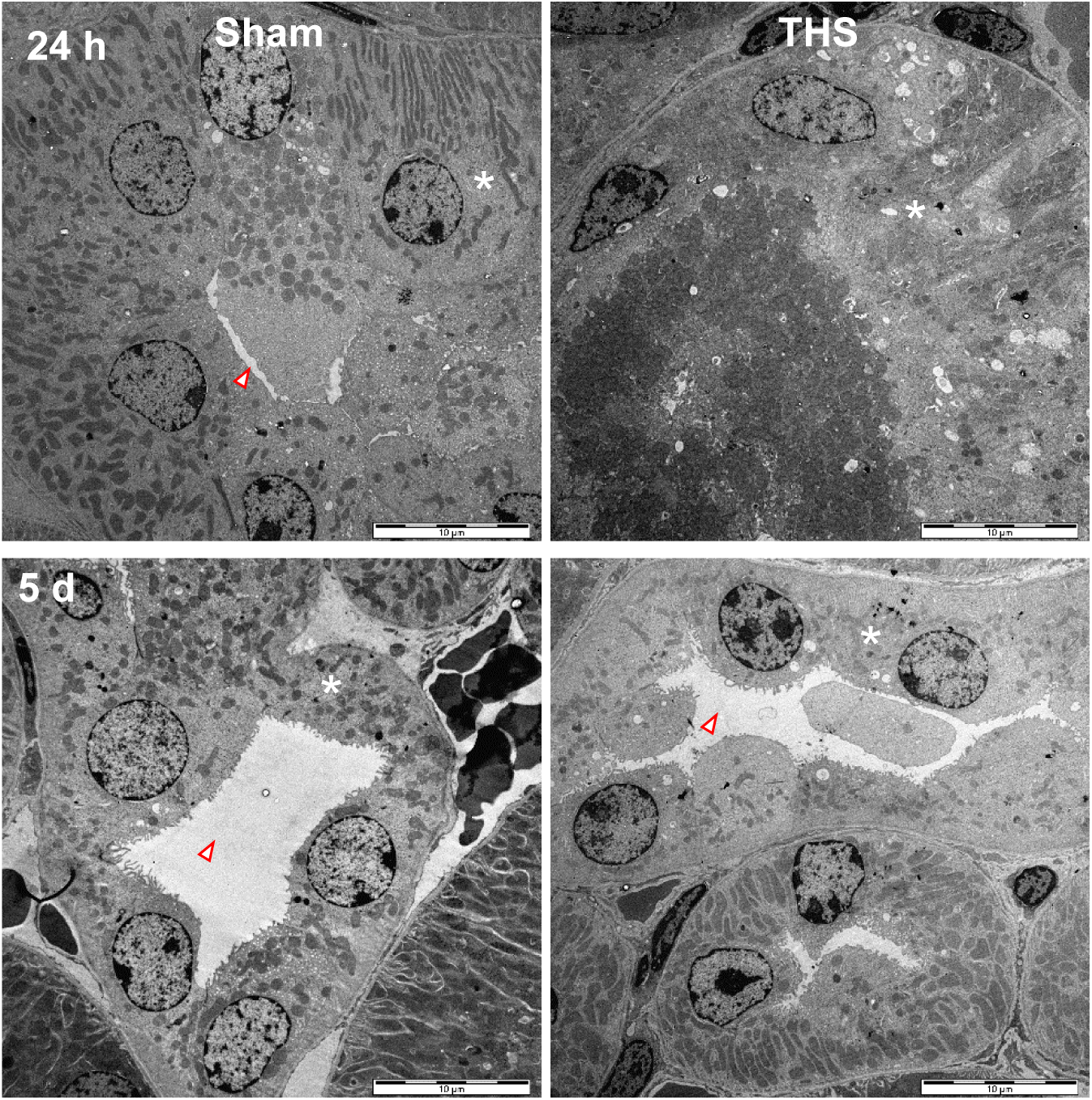
Transmission electron microscopy analysis of distal tubules. Kidneys 24 h and 5 d after trauma and hemorrhage (THS) or sham treatment were assessed using transmission electron microscopy. Representative images of distal tubules for *n* = 5-6 animals per group are shown. Star symbols indicate tubular cell cytoplasm; arrowheads indicate tubular lumen. Scale bars indicate 10 µm.

**Supplemental Figure 3.**
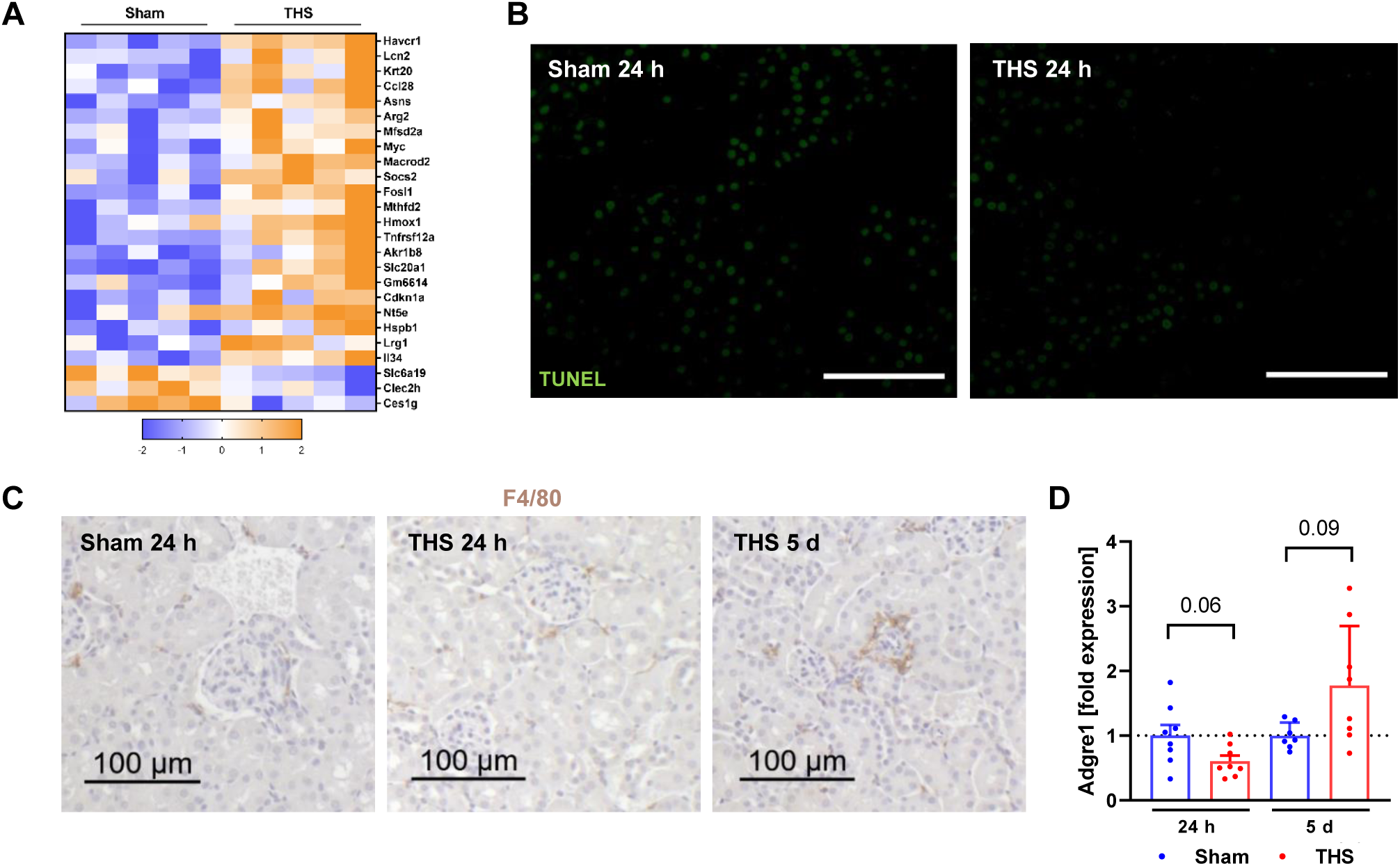
Renal overall gene expression, tubular apoptosis, and monocytes/macrophages. **A,** Whole-organ gene expression in kidneys from mice 24 h after THS or sham treatment (*n* = 5) was analyzed using a microarray. Significantly altered genes (*p* < 0.05 using Student‘s t-test, fold change ≥ 1.5) are shown. **B**, TUNEL staining in kidneys explanted 24 h post THS or sham treatment. Scale bars, 100 µm. **C**, F4/80 staining in kidneys at the indicated time points after THS or sham treatment. Monocytes/macrophages are stained in brown, nuclei in blue. Images are representative of at least *n* = 5 animals per group. **D**, Tissue expression of F4/80 (gene: Adgre1) 24 h and 5 d after THS or sham treatment.

**Supplemental Figure 4.**
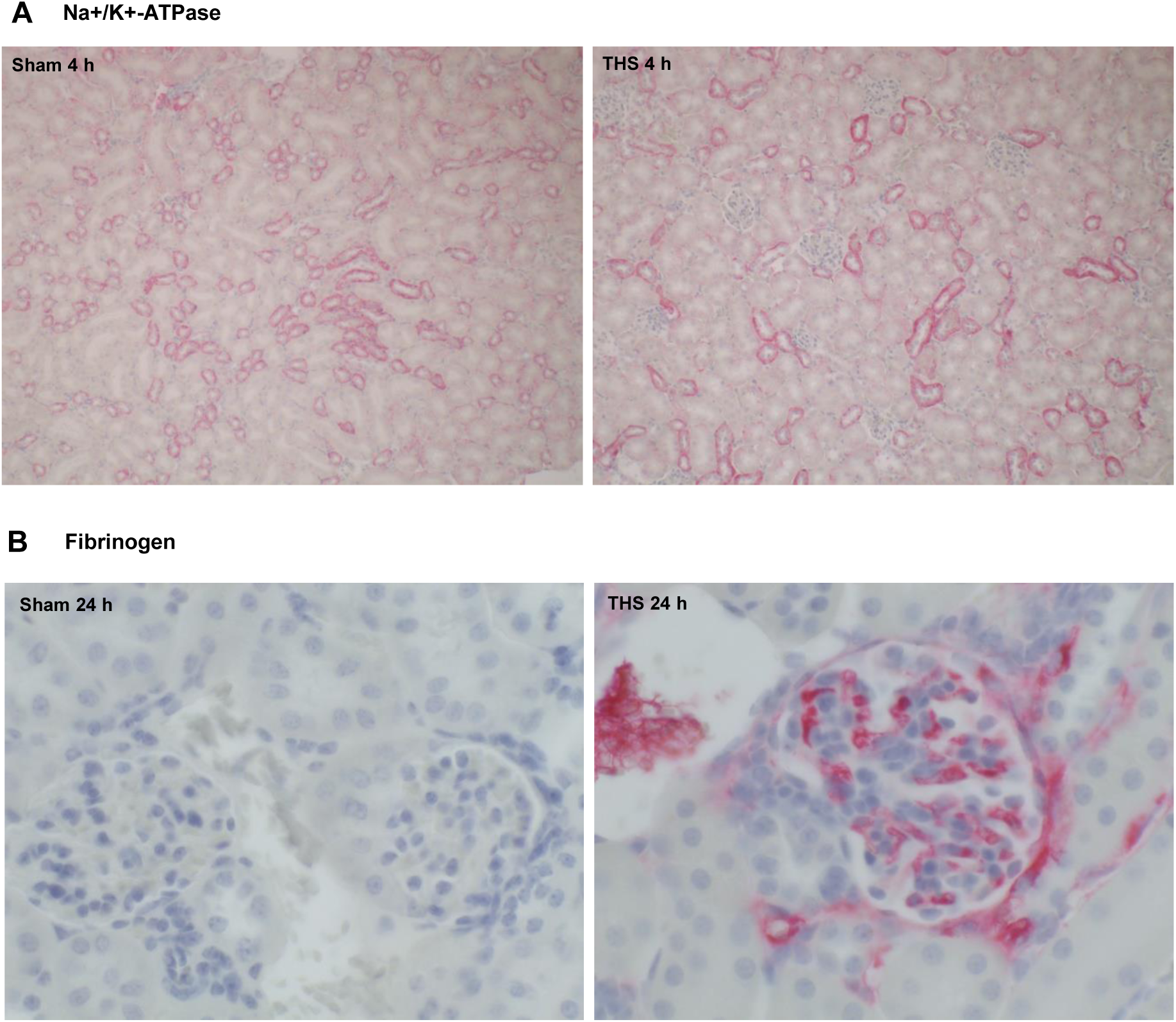
Tissue distribution of the Na^+^/K^+^-ATPase and of fibrinogen in kidneys after trauma. Representative images of the renal expression of **A**, Na+/K^+^-ATPase 4 h after THS or sham treatment and **B**, fibrinogen 24 h after THS or sham treatment as detected by immunohistochemistry (staining in red, nuclei in blue) of *n* = 4-8 animals per group. Magnification 100x (**A**) and 400x (**B**).

**Supplemental Table 1:**
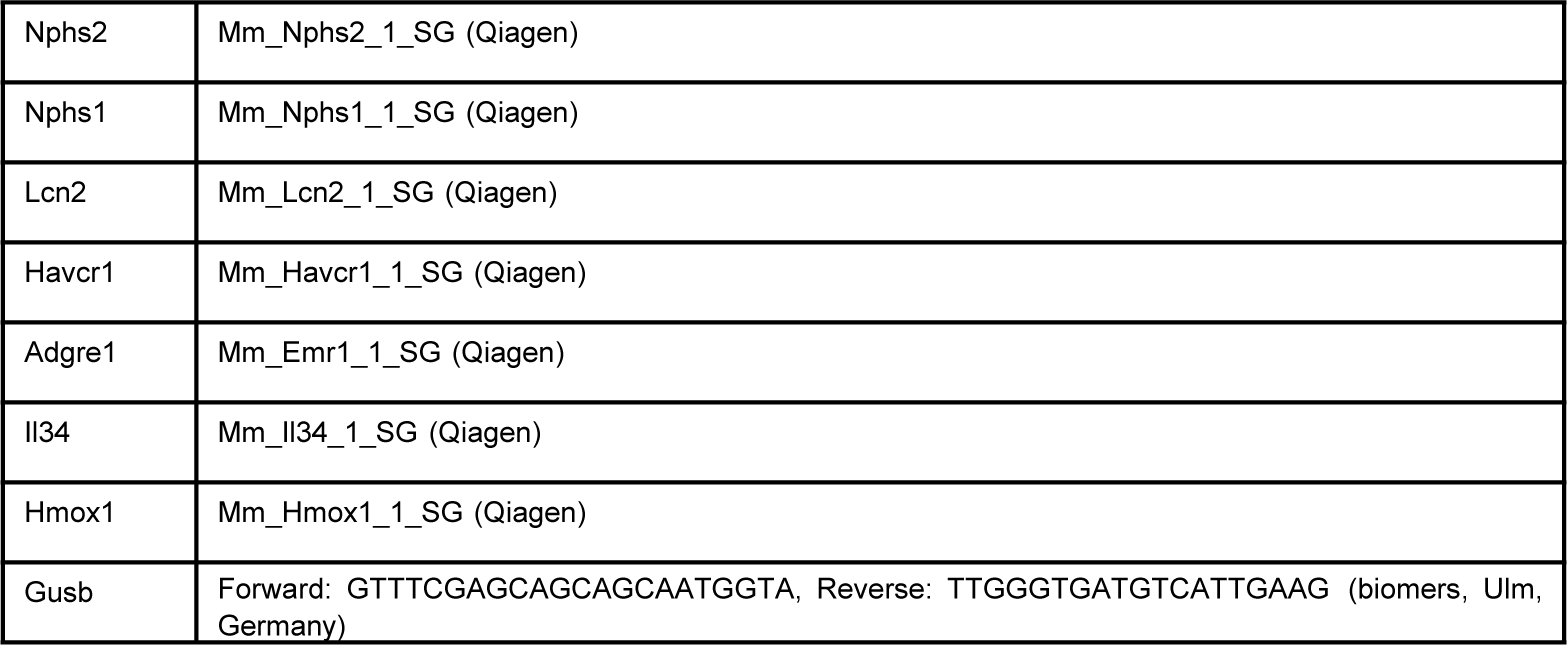
Primers used for qPCR analysis:

**Supplemental Table 2:** Settings during contrast-enhanced microfocused computed tomography.

| Parameter | Setting |
| --- | --- |
| Voxel size [µm] | 2 |
| Tube voltage [kV] | 90 |
| Tube current [µA] | 90 |
| Exposure [ms] | 750 |
| Average | 2 |
| Skip | 1 |
| Number of images | 2100 |
| Scan time [min] | 80 |

